# A national assessment of waterbird hunting in coastal wetlands of Suriname, South America

**DOI:** 10.1101/2025.04.04.646930

**Authors:** David S. Mizrahi, Arie L. Spaans, Marianne J. Spaans-Scheen, Amelia R. Cox, Benoit Laliberte, Eduardo Gallo-Cajiao, Christian Roy

## Abstract

The central northern coast of South America has extensive wetlands critical for waterbird conservation. While waterbird harvest occurs in the region, the impact on species population dynamics remains unclear. This study assesses waterbird hunting in the coastal wetlands of Suriname, addressing: (i) the extent of waterbird harvest, (ii) changes in harvest magnitude over time, and (iii) the motivations and methods used by hunters. We collected data via a national survey of licensed hunters in 2006 and 2016 using structured interviews. A Bayesian hierarchical model was used to analyze the data. We estimated harvest levels for 11 species and three groups (small herons, small shorebirds, large shorebirds). For most species, mean harvest per hunter significantly decreased from 2006 to 2016, except for blue-winged teal and migratory shorebirds. Most hunting was for non-commercial purposes (personal consumption and recreational). This is the first national assessment of waterbird hunting in Suriname. Harvest levels vary by species, and the sustainability of these levels remains uncertain. Managing hunting in Suriname requires addressing both legal and illegal hunting. Given Suriname’s importance for waterbirds, particularly species like the scarlet ibis and migratory shorebirds, it should be a priority for conservation efforts.

## 1. Introduction

Wetlands are biologically productive ecosystems that have sustained humans globally. The use of wetlands has included the harvesting of plants and animals, such as fish and waterbirds, with multiple purposes (e.g., cultural, subsistence, recreational, commercial). However, following increased pressure on the environment during the second phase of the Anthropocene (i.e., the Great Acceleration), the exploitation of some species may have contributed to population declines, and in some cases, extinction (Steffen et al. 2011; Benitez-Lopez et al. 2017; Cornford et al. 2023). Additionally, for long-distance migratory species, such as birds, harvest at specific locales can be compounded over large spatial scales and become unsustainable (e.g., Gallo-Cajiao et al. 2020). Consequently, it is necessary to evaluate harvest levels, their impacts on population dynamics, and the social and economic factors that drive them, in order to inform effective conservation and management policies for wetland-dependent species. Against this backdrop, there is a strong need to assess waterbird harvest worldwide, but particularly tropical regions, where people’s livelihoods often rely on natural resources, governance capacity for wildlife management may be weak, and harvest monitoring is relatively limited (Amano et al. 2018; Brotherton et al. 2020).

The central northern coastal region of South America include extensive wetlands crucial for waterbird species conservation (Butler et al. 2001; Murray et al. 2019). These ecosystems comprise mangrove forests, brackish lagoons, freshwater swamps, and the most extensive mudflats of South America, a flat coastline, and is strongly influenced by the Guiana Current. Noteworthy, this region has a low human footprint with little intertidal wetland losses (Murray et al. 2022), so threats associated with land use change are generally not prevalent (Allan et al. 2023). These coastal wetlands serve as important non-breeding and stopover areas for migratory shorebirds (i.e., Aves: Scolopacidae, Charadriidae) along the Western Atlantic Flyway. Additionally, over 100 species of waterbirds, many of which are non-migratory, occur in these coastal wetlands, including ducks (Anatidae), flamingoes (Phoenicopteridae), and herons (Ardeidae). Along the central northern coast of South America, Suriname has the largest concentration of some waterbird species (Spaans 1978; Morrison & Ross 1989), including scarlet ibis (*Eudocimus ruber*), semipalmated sandpiper (*Calidris pusilla*), greater yellowlegs *(Tringa melanoleuca*), lesser yellowlegs (*Tringa flavipes*), and short-billed dowitcher (*Limnodromus griseus*). The importance of this region has been recognized through the designation of Important Bird Areas by BirdLife International, Ramsar wetlands of international importance, and sites of hemispheric importance under the Western Hemisphere Shorebird Reserve Network (Ramsar Convention Secretariat 1997; Western Hemisphere Shorebird Reserve Network 2024; BirdLife International 2026).

Waterbird population trajectories in this region are generally unknown with some exceptions. For instance, the central northern coast of South America is not surveyed as part of the International Waterbird Census, hindering long-term monitoring of populations (Wetlands International 2026). This data gap has also impeded assessments of protected area effectiveness, which is essential for management (Wauchope et al. 2022). However, some species that occur in this region are declining across their entire range, such as the scarlet ibis, wood stork (*Mycteria americana*), and white-cheeked pintail (*Anas bahamensis*; BirdLife International 2025). Similarly, several migratory shorebirds species (e.g., semipalmated sandpiper, lesser yellowlegs, and short-billed dowitcher) have shown a significant decline of over three decades along the central northern coast of South America (Morrison et al. 2012), consistent with overall declines observed in North America (Smith et al. 2023).

Harvest, whether for commercial, recreational or subsistence purposes, can significantly contribute to mortality rates and influence population dynamics of some waterbird species. Hence, harvest assessments can play a crucial role in the development of sustainable harvest management frameworks and to inform conservation objectives (Watts et al. 2015; Koneff et al. 2017; Cox et al. 2024). Particularly, harvest assessments are important for shorebirds, as they typically have slow life history traits, such as small clutch sizes and only one breeding attempt per season, making them highly vulnerable to high harvest levels or even to seemingly low harvest levels when populations are declining (Watts et al. 2015). For migratory bird species, harvest assessments must also occur in multiple countries to account for the cumulative effects of harvest to be estimated. Recent studies from Guyana (Andres et al. 2022), French Guiana (Taylor 2017), and Brazil (Bosi de Almeida et al. 2018) show widespread shorebird hunting in the central northern coast of South America. Beyond this specific region, migratory shorebirds have also been hunted along the Western Atlantic Flyway in parts of the Lesser Antilles (Watts & Turrin 2016; Gebauer 2018; Reed et al. 2018). In the United States and Canada, the extent of hunting remains unclear, as part of subsistence hunting (but see Naves et al. 2019) and illegal harvest during recreational hunting (Katzner et al. 2020).

In addition to evaluating harvest from an ecological standpoint, understanding the human dimensions of hunting is crucial for informing conservation policy. These dimensions include parameters such as demographics of resource users (e.g., age, gender, income), hunting’s contribution to livelihoods, and attitudes toward hunting and regulations (Vaske & Manfredo 2012; Eriksson et al. 2023). For instance, hunting of migratory shorebirds in Southeast Asia can make a considerable contribution to household income (Zöckler et al. 2010; Gallo-Cajiao et al. 2020). In Guyana, shorebird hunters are typically fishermen supplementing their income when fishing conditions are poor (Andres et al. 2022). Understanding these parameters in the context of the central northern coast of South America could help inform more effective and equitable harvest management policies.

Here, we present a case study on a national assessment of waterbirds harvested in coastal wetlands of Suriname. Our aim is to contribute to the evaluation of harvest sustainability of waterbird species in Suriname within the context of central northern South America and the Western Atlantic Flyway. Specifically, we ask: (i) what is the recent extent of waterbird harvest in coastal wetlands of Suriname?; (ii) has magnitude of waterbird harvest changed over time?; and, (iii) what are the motivations of waterbird hunters and, what harvest methods they use?

## 2. Methods

### 2.1 Study area

Suriname, located on the central northern coast of South America, covers 163,800 km^2^ and has a 386 km coastline. It borders the Atlantic Ocean, Guyana, French Guiana, and Brazil. The country has a tropical climate with two dry and two wet seasons. Major rivers include the Courantyne, Coppename, Saramacca, Suriname, Commewijne, and Marowijne. Most of Suriname’s population (87%) lives in the coastal area, where the economy is primarily driven by mining and agriculture (Food and Agriculture Organization 2024). The coastal zone, mostly low and flat, comprises the lower and upper coastal plains, which are key waterbird habitats (Spaans et al. 2018). Wetland ecosystems in the lower coastal plain include mudflats, beaches, estuaries, mangroves, salt pans, coastal marshes, swamp forests, and shallow brackish lagoons. The upper coastal plain includes herbaceous and peat swamps. Our study focused on all coastal districts except Marowijne, where hunters were difficult to reach (Figure 1).

**Figure 1.**
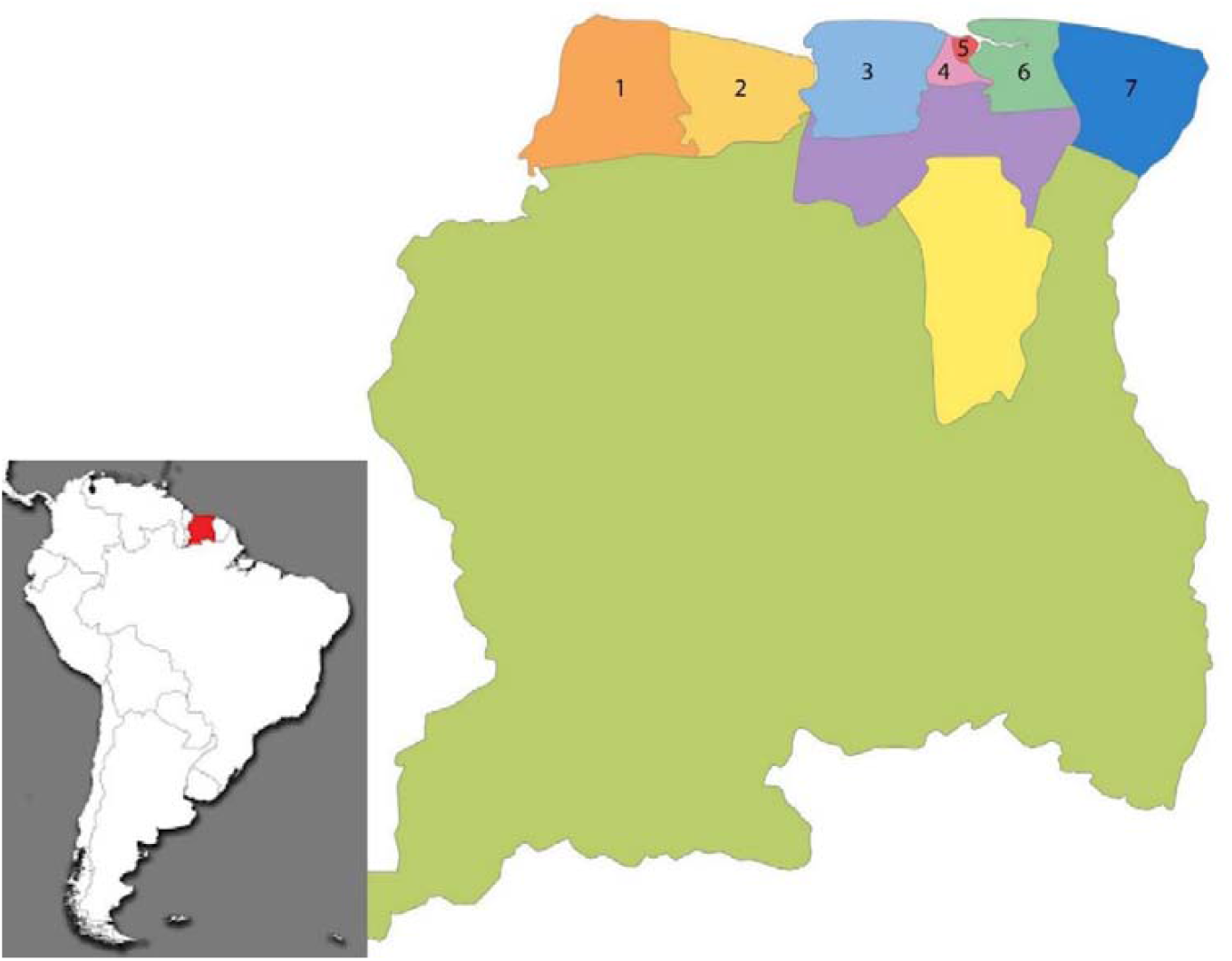
Map of Suriname with district borders, showing coastal districts numbered: 1) Nickerie, 2) Coronie, 3) Saramacca, 4) Wanica, 5) Paramaribo, 6) Commewijne, and 7) Marowijne. Note: the Marowijne district and non-coastal districts (Para [purple], Brokopondo [yellow] and Sipaliwini [green]) were not surveyed during this study.

The coastal zone of Suriname has various designations under different international institutional frameworks, including one wetland of international importance under the Ramsar Convention, three hemispherically important sites under the Western Hemisphere Shorebird Reserve Network, and four Important Bird Areas under BirdLife International (Ramsar Convention Secretariat 1997; Western Hemisphere Shorebird Reserve Network 2024; BirdLife International 2026). Domestically, there are four Multiple-Use Management Areas (MUMAS) and four nature reserves as well (Spaans et al. 2018). Some of these areas have overlapping designations.

The Hunting Act 1954 and the corresponding implementing legislation (i.e. Hunting Decree 2002 as amended in 2009), determine the species of wildlife that may be hunted and when. All bird species occurring in Suriname are fully protected, except those designated as ‘game’, ‘cage birds’, and ‘predominantly harmful species’. Waterbird species without full protection under Suriname regulations and considered in this paper are: black-bellied whistling-duck (*Dendrocygna autumnalis*), muscovy duck (*Cairina moschata*), white-cheeked pintail, blue-winged teal (*Spatula discors*), Neotropic cormorant (*Nannopterum brasilianum*), and anhinga (*Anhinga anhinga*; Table 1).

**Table 1:**
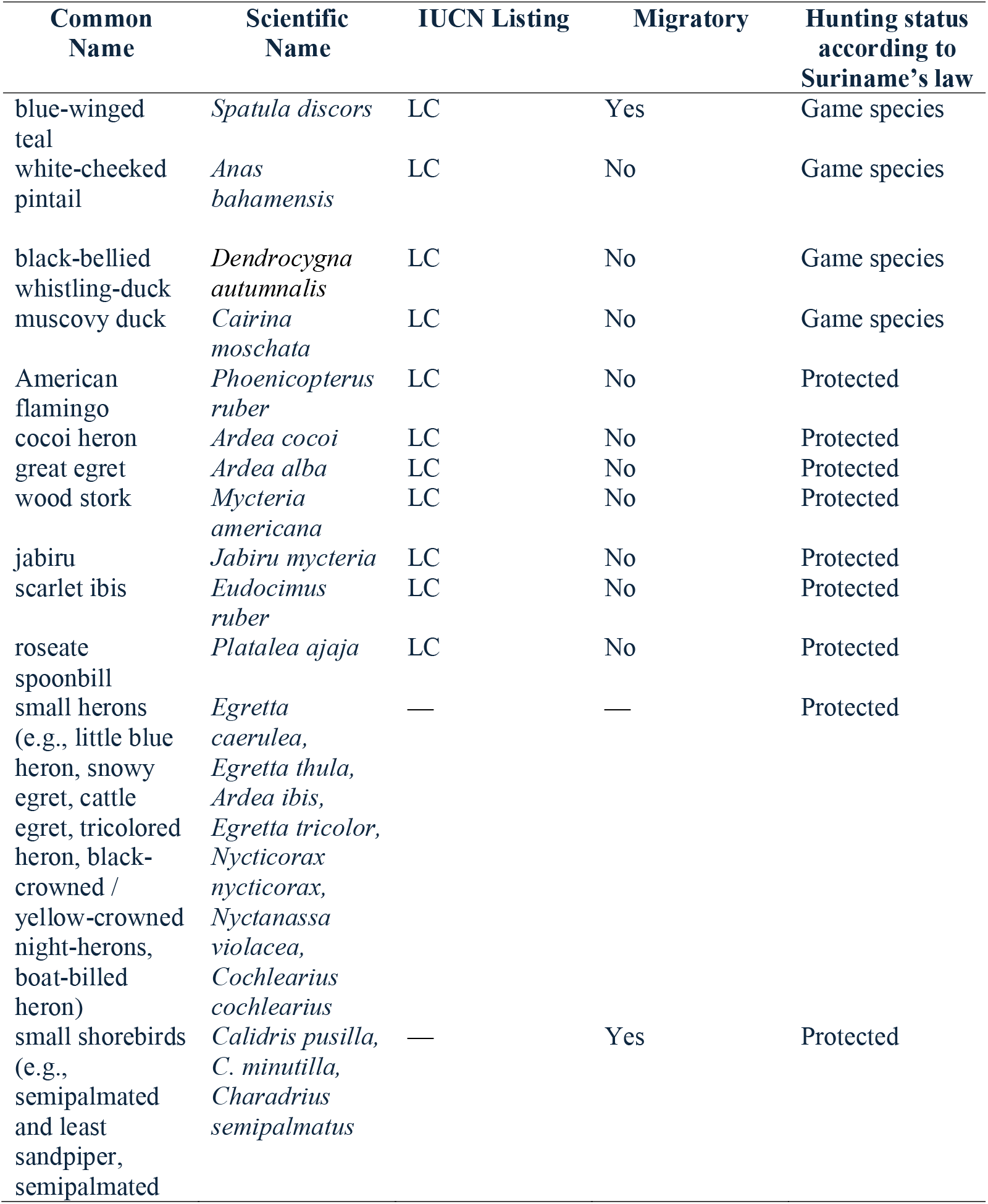
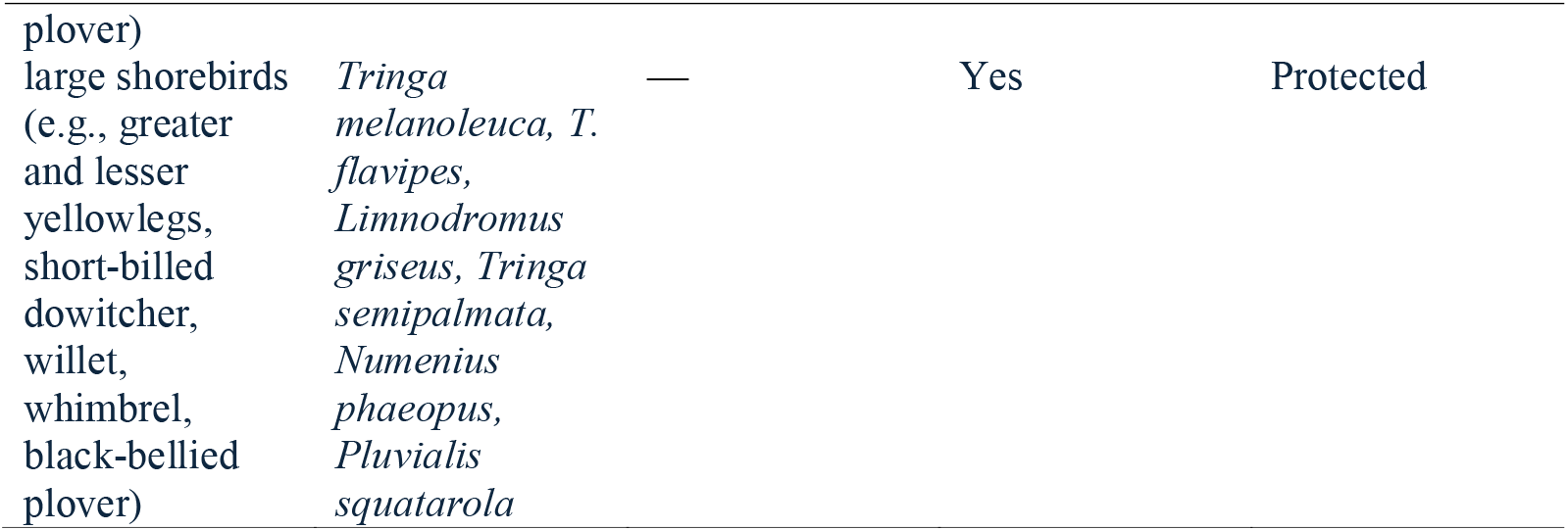
Species and species groups of waterbirds included in the hunter survey conducted in six districts of Suriname in 2006 and 2016. International Union for Conservation of Nature’s (IUCN) Red List of Threatened Species information was sourced from the version 2022-2 (IUCN 2022).

| Common Name | Scientific Name | IUCN Listing | Migratory | Hunting status according to Suriname's law |
| --- | --- | --- | --- | --- |
| blue-winged teal | *Spatula discors* | LC | Yes | Game species |
| white-cheeked pintail | *Anas bahamensis* | LC | No | Game species |
| black-bellied whistling-duck | *Dendrocygna autumnalis* | LC | No | Game species |
| muscovy duck | *Cairina moschata* | LC | No | Game species |
| American flamingo | *Phoenicopterus ruber* | LC | No | Protected |
| cocoi heron | *Ardea cocoi* | LC | No | Protected |
| great egret | *Ardea alba* | LC | No | Protected |
| wood stork | *Mycteria americana* | LC | No | Protected |
| jabiru | *Jabiru mycteria* | LC | No | Protected |
| scarlet ibis | *Eudocimus ruber* | LC | No | Protected |
| roseate spoonbill | *Platalea ajaja* | LC | No | Protected |
| small herons (e.g., little blue heron, snowy egret, cattle egret, tricolored heron, black-crowned / yellow-crowned night-herons, boat-billed heron) | *Egretta caerulea*,<br>*Egretta thula*,<br>*Ardea ibis*,<br>*Egretta tricolor*,<br>*Nycticorax nycticorax*,<br>*Nyctanassa violacea*,<br>*Cochlearius cochlearius* | — | — | Protected |
| small shorebirds (e.g., semipalmated and least sandpiper, semipalmated) | *Calidris pusilla*,<br>*C. minutilla*,<br>*Charadrius semipalmatus* | — | Yes | Protected |
| plover) |  |  |  |  |
| large shorebirds | *Tringa* | — | Yes | Protected |
| (e.g., greater | *melanoleuca*, *T.* |  |  |  |
| and lesser | *flavipes*, |  |  |  |
| yellowlegs, | *Limnodromus* |  |  |  |
| short-billed | *griseus*, *Tringa* |  |  |  |
| dowitcher, | *semipalmata*, |  |  |  |
| willet, | *Numenius* |  |  |  |
| whimbrel, | *phaeopus*, |  |  |  |
| black-bellied | *Pluvialis* |  |  |  |
| plover) | *squatarola* |  |  |  |

### 2.1 Study design and data collection

This is a cross-sectional and longitudinal single case study based on surveys conducted at two points in time that are ten years apart. Travel is difficult, particularly on unpaved roads during the rainy season in Suriname, so data were collected in April 2006 and 2016, during what is typically the short dry season, to improve our chances of reaching hunters in their communities.

At the time the surveys were conducted, no ethics approval was required in Suriname. The surveys were conducted in collaboration with the Suriname Forest Service/Nature Conservation Division, a department within the Ministry of Land Policy and Forest Management. Staff from the agency were actively involved in survey implementation.

In 2016, potential participants were identified from hunter licensing records provided by the District Commissioners in each of the included districts. Neither the District Commissioners nor their staff were informed of the identities of the selected participants. No minors were involved in the study, as individuals under the age of 18 are not permitted to hold a hunting license in Suriname. The participant pool consisted of those who could be reached and agreed to participate. Due to challenges in contacting hunters, we could not create a stratified sample proportional to the number of hunters in each district.

Prior to administering the survey, all participants were informed of the study’s purpose and how their responses would be used. They were advised that submission of a completed survey would be considered as informed consent to participate. While participants were encouraged to answer all questions, they were also clearly informed that responding to any specific question was entirely voluntary. All surveys were administered in Dutch, the official language of Suriname. Survey responses were recorded on paper and later digitized. All responses remained anonymous; no names or identifying information were included on the questionnaires. Because we did not record personal identification information, we could not intentionally interview any hunters from the 2006 survey in 2016.

The survey questions in 2006 focused primarily on hunting frequency, magnitude and purpose, and species or group of species of waterbirds harvested (Appendix 1). For some taxa, species reporting was not possible, or at least not accurately as hunters do not distinguish between species, so for these we present data for groups of species which share broad morphological and ecological traits (Table 1). The 2016 survey used all the same questions used in 2006 but included additional questions to improve our understanding of the hunter community (e.g., age, family members, hunting purpose), hunting seasonality, and hunting method (Appendix 2).

Survey respondents self-reported their harvest, a common method for assessing waterbird hunting (Geissler 1990; Smith et al. 2022). However, this approach has potential limitations, including recruitment bias and underreporting of mortality, such as wounded birds which may die later (i.e., crippling losses). In 2006, we piloted group surveys with hunters, but found that participants were conferring with each other, making the results unreliable. As a result, we switched to individual one-on-one interviews at hunters’ homes, continuing this method in 2016. If interviewers suspected dishonesty, they cross-checked answers with game wardens familiar with the hunters’ activities. Data deemed unreliable were excluded from analyses. All data were kept anonymous and confidential, and participation was voluntary.

### 2.2 Data analysis

#### 2.2.1 Estimation of licensed hunters in 2016

In 2006, there was no reliable centralized hunter license database, so the total number of licensed hunters that year is unknown. For 2016, we used the Suriname hunter license database, which tracks only license registrations. According to Suriname law, hunting licenses are valid for up to five years, from January 1st to December 30th. We considered the total number of licensed hunters in 2016 as the unique registrations between January 1, 2011, and December 30, 2016. This method does not account for licenses that may have been suspended before 2016, as the database lacked records on renewals or suspensions, and no corrective measures were applied.

#### 2.2.2 Estimation of hunting levels

We used a Bayesian hierarchical model to analyze the harvest survey data (Appendix 3). We modelled harvest data for 11 waterbirds species and four taxa (i.e., small herons, small shorebirds, large shorebirds, and all shorebirds). Because of the limited number of respondents in each region, we pooled all the data together and did not consider variation in hunting across regions/districts. In 2006, survey respondents were asked to provide estimates for all shorebird species combined. In 2016, respondents were asked to distinguish between harvest of large and small shorebird species. Therefore, the survey of small and large shorebirds was set to zero for 2006 and we modeled the harvest of all shorebirds directly. For 2016, we instead modelled the harvest of small and large shorebirds and derived the harvest of all shorebirds from those two categories.

During the interview process, hunters reported the number of birds they harvested that year via 8 categories (either 0; 1; 2 – 10; 11 – 50; 51 – 100; 101 – 500; 501 – 1000; or > 1000 birds). We therefore modeled the reported harvest of each species *i* by each individual hunter interviewed *j* via a categorical distribution:

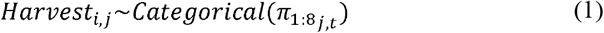

where *π*_1:8_ represents the vector of probabilities that the hunter will report their harvest in that category for a given species *j* in year *t*.

We assumed that the underlying distribution of harvest of each species would follow a negative binomial distribution, which allows for over dispersion that we observed in harvest data. We parametrized the negative binomial distribution via the mean and variance (Lindén & Mäntyniemi 2011) and modelled the mean harvest (*µ*_*j*,*t*_) for a given species *j* in year *t* such as:

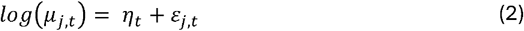

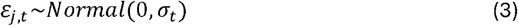

where *η*_*t*_ is the average harvest across all species on the log scale and a *ε*_*t*,*j*_ is species specific derivation from this mean in this given year. The species specific derivations are drawn from a normal distribution with year specific standard distribution. This hierarchical formulation allowed us to share information across species (i.e. ‘borrowing the strength’ from the ensemble) but to also consider the change in harvest level across years. We then parametrized the variance 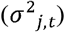 of the model in function of the mean such as:

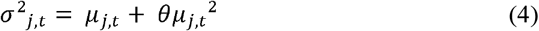

where *θ* is the overdispersion parameter for the quadratic mean–variance relationship of the classic negative binomial regression (Lindén & Mäntyniemi 2011). We used moment matching to link estimates of the mean and variance to the species-specific and year specific probability 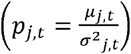 and size parameter 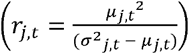 and then linked the probability of the harvest categories with the cumulative probability distribution (CDF; *F*(*k, p*_*j*,*t*_,*r*_*j*_)) of the negative binomial distribution. The first category of harvest represents the probability that harvest for a given species each year is exactly zero and can be derived directly from the CDF:

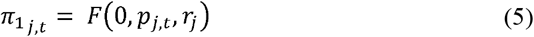

The probability of the harvest for categories 2 to 7 can be derived by subtracting the CDF at the lower bound of the category from the CDF at the upper bound of category. For example, for the probability that harvest is between 2 to 10 birds (i.e., Category 3) for a given species in a given year can be estimated via:

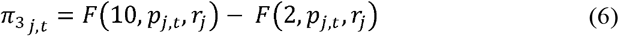

The last harvest category (*π*_8_;> 1000 *birds*)is the probability that the harvest is larger than the lower bound of the category:

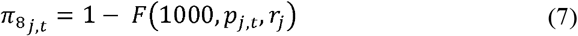

Hunters did not always complete the survey and did not report their harvest for all the species. In those cases, we used a data augmentation scheme and let the model assign a harvest category for the hunter. Because of the hierarchical formulation of the model, the category of harvest assigned to this hunter by the model will tend to be close to the most frequent category of harvest observed for that species during that year.

Finally, we derived the total annual harvest in 2016 by multiplying the number of registered hunters with the mean annual harvest (*µ*_*j*,2016_) for a given species *j*. Since the Suriname hunter license database was incomplete for 2006, we were not able to estimate the number of registered hunters and total annual harvest in 2006.

We used R version 4.2.0 for data processing (R Core Team 2020) and JAGS version 4.3.0 (Plummer 2003), called via the R package jagsUI (Kellner 2021) for analyses. We used non-informative priors for all parameters and ran three chains with randomized initial start values for 69,500 iterations, using the first 2,000 iterations as burn in and thinning the chains to save every 12th iteration, with 1,000 iterations of adaption yielding 45,000 posterior samples for each parameter. We assessed model convergence visually and by making sure that the Gelman-Rubin statistic 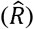 was less or equal than 1.1 (Gelman et al. 2013). We also used a posterior predictive check to evaluate the performance of the model. All results are reported as the median with 90% Bayesian Credible Intervals, calculated as high-density intervals using *bayestestR* package (Makowski et al. 2019)

## 3. Results

### 3.1 Hunters in Suriname

Surveys were conducted in six districts in 2006 and five districts in 2016, with the highest number of participants from Nickerie, Saramacca, and Commewijne (Table 2). Additionally, we were unable to recruit the same number of hunters from each district over the two time periods. Paramaribo District, despite having the largest population of licensed hunters, had the fewest participants (Table 2).

**Table 2:** Numbers of hunters interviewed about their harvest of waterbirds per district in Suriname in 2006 and 2016.

| District | 2006 | 2016 |
| --- | --- | --- |
| Nickerie | 115 | 186 |
| Coronie | 8 | 21 |
| Saramacca | 100 | 29 |
| Wanica | 3 | 11 |
| Paramaribo | 3 | 0 |
| Commewijne | 47 | 42 |
| **Total** | **276** | **289** |

In 2006, 276 licensed hunters answered the survey (Table 2) but we excluded 45 of them from the analysis as the answers were deemed to be unreliable. Of the remaining 231 surveys, only 69% (n=159) of the hunters reported their harvest of all species included in the survey. Hunters completed the survey and reported their harvest most frequently for muscovy duck (91%), black-bellied whistling-duck (91%), white-cheeked pintail (90%), while response rates were lower for jabiru (*Jabiru mycteria*; 82%) and American flamingo (*Phoenicopterus ruber*; 81%).

In 2016, a total of 289 male hunters answered the survey (Table 2) out of the 4727 hunters that were registered in the Suriname hunter license database. The answers to 16 surveys were deemed to be unreliable so we kept 273 surveys for the analysis. The ages of the respondents ranged from 19 to 85 years, with a mean age of 49 (SD = 11). Of the 273 surveys 99% (n=270) of the hunters reported their harvest for all species included in the survey. Of the surveyed hunters, 190 hunters indicated that they did not harvest any birds, 72 hunters reported harvesting 5 species or less, 8 hunters harvested between 6 and 10 species, and 3 hunters harvested more than 10 species.

In addition to the 11 species and four taxa that we modelled, hunters also reported the harvest of Neotropic cormorant, anhinga, brown pelican (*Pelecanus occidentalis*), black skimmer (*Rynchops niger*), and undefined Laridae taxa (i.e., *gulls and terns*). However, because the harvest levels for these species were too low for statistical modeling, we excluded them from the analysis.

### 3.2 Estimated waterbirds harvest in Suriname

#### 3.2.1 Average harvest for 2006 and 2016

In both 2006 and 2016, mean harvest per hunter of all shorebirds was higher than for any other species or group (Figure 2). Results from the 2016 survey indicate that hunters harvested on average more small shorebirds than large shorebirds. After shorebirds, the most harvested species in 2006 were black-bellied whistling-duck, scarlet ibis, and cocoi heron; the jabiru and American flamingo were the least harvested species (Figure 2). In 2016, the most harvested species outside of shorebirds was blue-winged teal followed by black-bellied whistling-duck, while American flamingo and the jabiru were still the two least harvested species (Figure 2).

**Figure 2:**
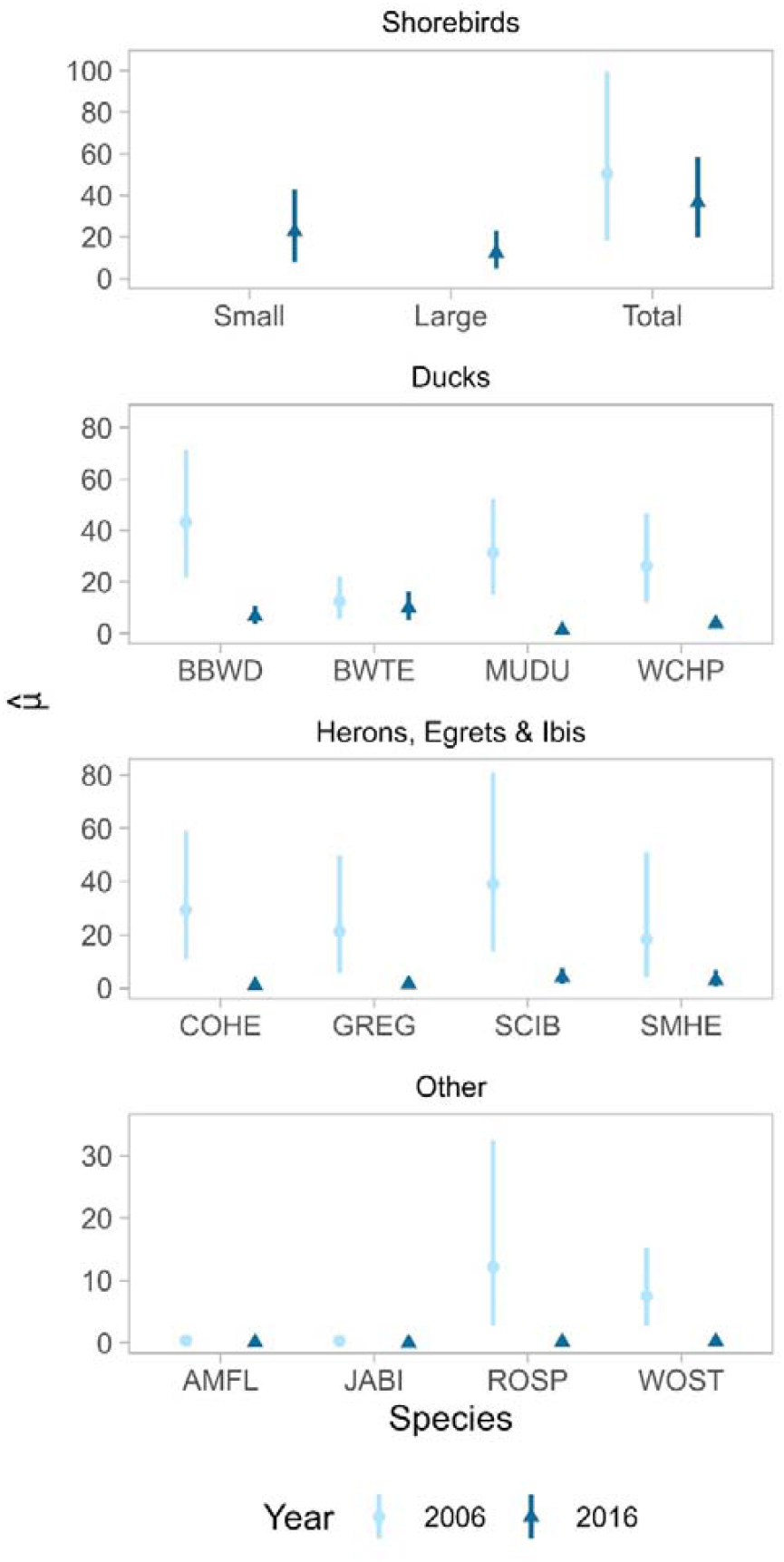
Mean birds harvested in a year (μ) per hunter based on surveys of hunters in Suriname in 2006 and 2016. Data points represent the median, with error bars indicating the 90% high-density intervals. American Ornithological Society species codes are presented in Table 3. Note that in 2006, hunters reported all shorebirds species combined, but in 2016 large and small shorebirds were distinguished.

#### 3.2.2. Has hunting of waterbird species changed over time?

For most waterbird species, mean harvest per hunter declined between 2006 and 2016 (Figure 2). The exceptions were the blue-winged teal and the total migratory shorebirds (i.e., small and large combined), for which harvest did not decrease significantly between 2006 and 2016. In particular, wood stork, roseate spoonbill (*Platalea ajaja*), and jabiru harvest declined below one bird per hunter on average in 2016. Hunters surveyed in 2016 generally agreed that their overall harvest had declined. Among respondents who answered this question and hunted in both 2006 and 2016, most reported hunting less in 2016 (77%), while a small minority (7%) reported hunting more than in 2006. When asked why they had changed their hunting practices, out of 66 responses provided the most common response was that there were fewer birds available for harvest than there were in the past (29%). Some individuals also cited the increase in hunting regulations and increased enforcement (12%) and associated costs (6%).

**Table 3:**
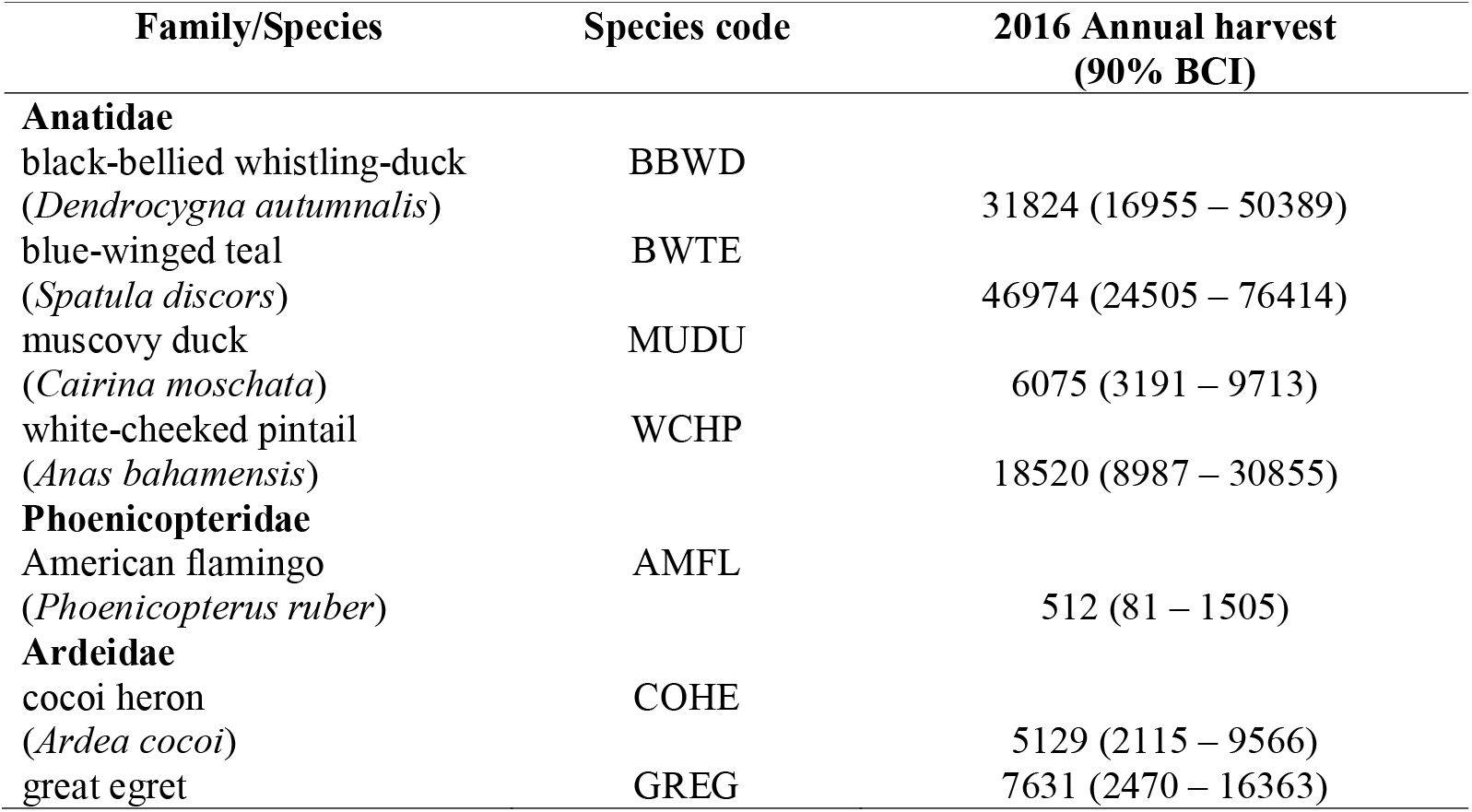
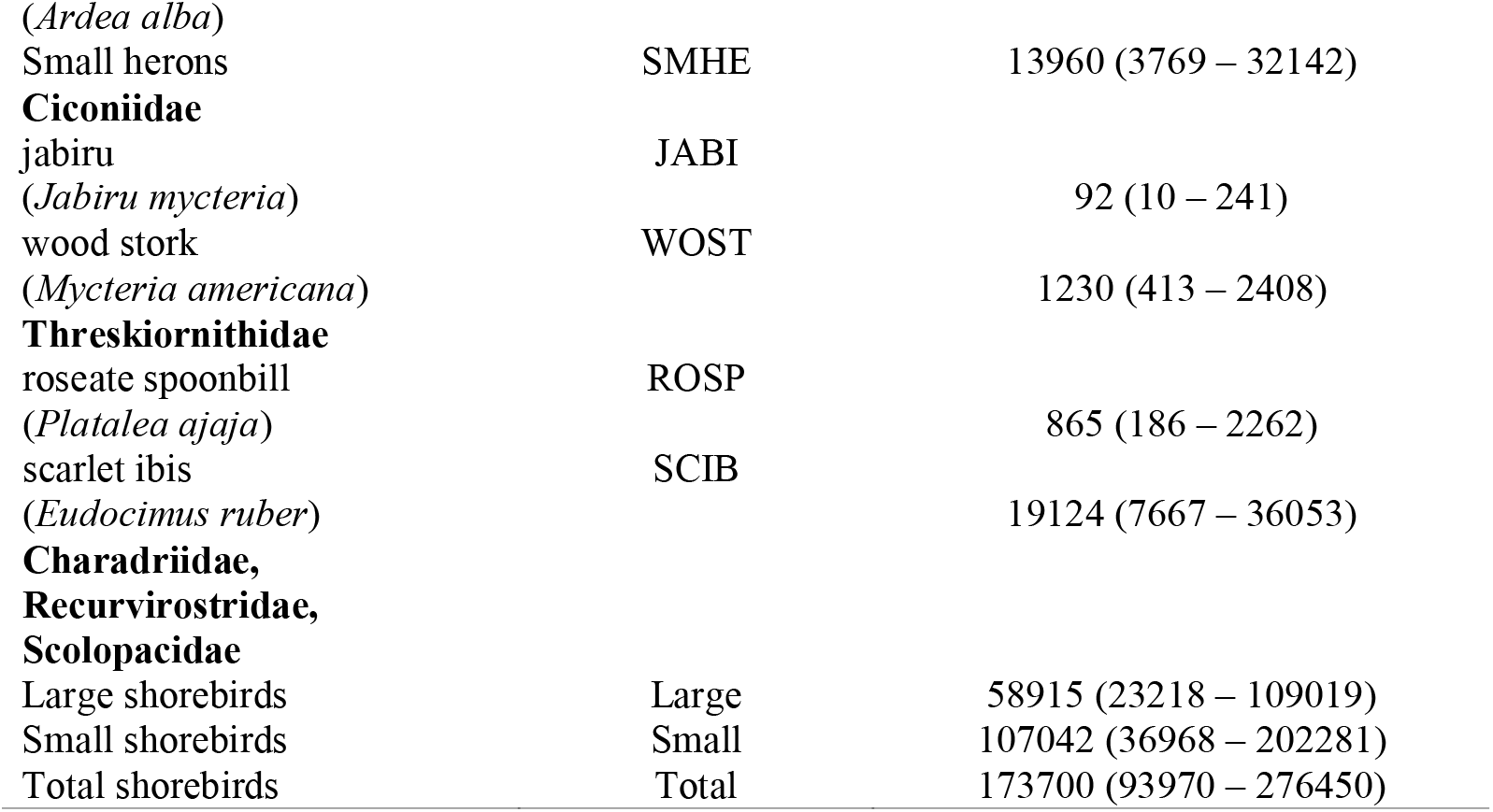
Estimated total waterbird harvest by all registered hunters in Suriname in 2016. Estimates are presented as medians with 90% Bayesian Credible Intervals (BCI).

#### 3.2.3. Total derived harvest 2016

Shorebirds were the most harvested group in general in 2016 with a total harvest of approximately 174 thousand birds (90% BCI = 94 – 276 thousand; Table 3). Small shorebirds (107 [90% BCI = 37 – 202] thousand) were harvested nearly twice as much as large shorebirds (59 [90% BCI = 23 – 109] thousand). Ducks were the next group of species with the most harvest, with the blue-winged teal having the highest total harvest (47 [90% BCI = 25 – 76] thousand), followed by herons, egrets and ibis, with the scarlet ibis (19 [90% BCI = 8 – 36] thousand; Table 3) having the highest total harvest. Finally, the wood stork, roseate spoonbill, American flamingo, and jaribu were less frequently harvested species with only the wood stork (1230 [90% BCI = 413 – 2408]; Table 3) harvest exceeding a thousand birds.

#### 3.2.4. What are the motivations of waterbird hunters and what hunting methods do they use?

Across both survey years (2006 and 2016), 63% of respondents travelled to their hunting location with the intention of engaging in another activity, generally fishing or recreation. Based on data from 2016 only, nearly all hunters who reported their hunting method (N=90) reported hunting primarily with a firearm (99%), while a small proportion reported using a net (1 %). In 2016, hunters who reported their primary purpose for hunting (N=90) indicated that it was predominantly non-commercial in purpose, which includes personal consumption and recreational activities. Though there is overlap between these two motivations, the majority of hunters reported hunting for the former (70%) and only a small proportion for the latter (13%). Additionally, 13% of the hunters reported selling the birds inferring a commercial purpose.

## 4. Discussion

Harvest of waterbird species is pervasive across coastal wetlands of Suriname, has declined over time, and is primarily practiced for personal consumption. To the best of our knowledge, this is the first national assessment of waterbird harvest from Suriname, part of a region of critical importance to shorebird conservation in central northern South America and along the Western Atlantic Flyway. While the importance of waterbird harvest in Suriname has been internationally recognized for many years (Ottema & Spaans 2008; AFSI Harvest Working Group 2017, 2020), no structured assessment has been previously conducted. Waterbird harvest occurs along the entire coast of Suriname and includes species across a wide range of taxonomic groups. Amongst all waterbirds hunted in Suriname, the scarlet ibis and migratory shorebirds are of particular conservation concern due to higher hunting levels, declining populations, and slow life history traits (Olmos & Silva e Silva 2001; Watts et al. 2015; BirdLife International 2016; Smith et al. 2023).

In Suriname, waterbirds are primarily harvested using firearms, with only a small proportion using alternative methods, such as nets and choking wires. This pattern is similar to French Guiana (Taylor 2017) but contrasts with neighboring Guyana and Brazil, where choking wires, fishing nets and night lighting are the most common methods (Bosi de Almeida et al. 2018; Andres et al. 2022). In line with broader understanding of wetland use by humans, waterbird harvest did not happen in isolation in our study but rather alongside other resource uses, particularly fishing.

Based on hunters’ self-reported motivations, waterbird harvest in Suriname was primarily non-commercial and driven by recreational interest and personal consumption; the latter was the most frequently cited motivation in our study. This pattern contrasts with Guyana, where commercial harvest of migratory shorebirds occurs at a relatively larger scale although the overall harvest is less than in Suriname (Andres et al. 2022). While the limited commercialization reported in Suriname is encouraging, cross-border transport of waterbirds from western Suriname to markets in Guyana has been documented and warrants further investigation. Beyond subsistence, some of the harvest done for personal consumption may have cultural dimensions, as suggested by the consumption of birds such as the scarlet ibis at social celebrations (Ottema & Spaans 2008), a practice noted elsewhere in the region as being driven by the bird’s status as a traditional delicacy (BirdsCaribbean 2017).

Species hunted represented a wide range of taxonomic groups and harvest levels varied between species, a pattern consistent with tropical wetlands elsewhere (Iskandar & Karlina 2004; Ramachandran et al. 2017; Bosi de Almeida et al. 2018; Deniau et al. 2022). Ducks were among the most heavily harvested groups,likely reflecting both their high availability and hunter preferences for larger-bodied species. Of all the duck species hunted, only the blue-winged teal is considered migratory. Harvest levels of blue-winged teal from Suriname were modest compared to North America, representing about 1.68% of the combined harvest estimated for the US and Canada during the same year (Raftovich et al. 2018). The scarlet ibis was also among the species with the highest levels of hunting, and although it is hunted elsewhere in the tropics, the levels reported here are unusually high compared to recent reports from other regions (Deniau et al. 2022). Harvest of other species of waterbirds such as the American flamingo, the wood stork, the jabiru, and the roseate spoonbill, were relatively low and in line with level reported for other areas such as the Sahel (Deniau et al. 2022).

Current estimates of migratory shorebird harvest in Suriname exceed those available for any other country along the central northern coast of South America and the Lesser Antilles. We estimated that ≈ 174 thousand migratory shorebirds were harvested in 2016, but this is a conservative estimate as it does not account for crippling loss, under reporting because of illegality or unlicensed hunters. While empirical data on crippling rates for shorebirds, and particularly in South America, are virtually non-existent, estimates from other migratory game birds provide valuable context. For species such as doves and waterfowl, crippling losses typically range from 10% to over 25% and may be influenced by various hunting factors, including the type of ammunition used (i.e., steel shot vs lead shot; Schulz et al. 2013; Ellis & Miller 2022; Moreno-Zárate et al. 2023). Nevertheless, our estimate for shorebird harvest in Suriname greatly exceeds combined estimates reported for Guyana, the Lesser Antilles, and Brazil (Bosi de Almeida et al. 2018; Andres et al. 2022; AFSI Harvest Working Group 2020).

Our findings are limited by the qualitative, self-reported nature of the survey data, including recall bias, rounding, or intentional misreporting. Broad harvest categories and assumptions about the underlying distribution may affect precision but are unlikely to alter the overall magnitude of our estimates. Misclassification near range category boundaries and limited information on extreme values may affect estimation of the underlying distribution, and the assumption of a negative binomial distribution may not fully capture its shape, potentially biasing mean and total harvest estimates. This may affect precision but are unlikely to alter the overall magnitude of our estimates. We also assumed surveyed hunters were representative of all registered hunters. Future surveys could improve accuracy through narrower harvest categories and a stratified sampling.

The sustainability of waterbird harvest in Suriname remains largely unknown, but inferences can be drawn from ecological theory and species life-history traits. The population-level effects of harvest range from compensatory to additive, or even super-additive mortality, depending on hunting intensity and species life history (Anderson & Burnham 1976). Additive mortality occurs when harvest reduces survival, while compensatory mortality arises when harvest does not affect survival due to density dependence or targeting individuals that would have died naturally (Sandercock et al. 2011; Cooch et al. 2014). Generally, species with large clutch sizes, high natural mortality, and large populations experiencing density dependence can generally sustain higher harvest rates than species with small clutch sizes, high annual survival, and small populations (e.g., most shorebird species; Nichols et al. 1984; Cooch et al. 2014). The timing of harvest relative to natural mortality is also important, with hunting timed during or after periods of high natural mortality more likely to be additive (Kokko 2001; Blomberg 2015). For many species, migration represents a period of elevated natural mortality particularly for juveniles (Lok et al. 2015; Rushing et al. 2017; Sergio et al. 2019).

While duck hunting may pose relatively lower conservation concern, uncertainty around population status of resident ducks in Suriname suggests a precautionary approach is warranted. The scarlet ibis population is declining across its range (Sainz-Borgo 2025), and given its slow life history traits (i.e., small clutch size, delayed sexual maturity; Olmos & Silva e Silva 2001), current harvest levels may be unsustainable. A more reliable assessment of the scarlet ibis population could encourage better adherence to regulations, as hunters may use uncertainty as a justification to dismiss the need for the law (i.e., neutralization theory; Eliason 1999). Compliance promotion campaigns could help, but poachers often comply only in the presence of law enforcement (Jacoby 2014; von Essen et al. 2014). Any harvest management strategies and compliance promotion efforts will need to consider broader socio-economic contexts such as local wetland use, livelihoods, and governance capacity to be effective (Ostrom 2009; Sutherland et al. 2014).

For migratory shorebirds, species-level harvest estimates are lacking, but concern is particularly high for the semipalmated sandpiper, a Near Threatened species (BirdLife International 2024) that likely accounts for a large percentage of the small shorebirds harvested in Suriname given its prevalence in the region (Morrison & Ross 1989). Shorebird harvest in Suriname remains high, and up to 44% of the flyway-wide sustainable hunting levels could already be taken from neighboring Guyana (Watts et al. 2015; Andres et al. 2022).

Most species in our study are protected and the harvest of flamingoes, herons, storks, ibises, and shorebirds in our surveys were illegal under Suriname’s Hunting Act (1954). The observed declines for the harvest of flamingoes, herons, storks, and ibises are encouraging. These declines are also concurrent with an increase in compliance promotion and enforcement of harvest regulations, which likely contributed to the reduction in take. Shorebirds harvest level remains problematic. The status of shorebirds as protected species also complicates the engagement of hunters over species identification and creates challenges for the assessment of shorebird harvest sustainability (Solomon et al. 2015). This situation is not unique to Suriname however and has been observed elsewhere such as in Guyana (Andres et al. 2022) and the East Asian-Australasian Flyway (Gallo-Cajiao et al. 2020).

Furthermore, a patchwork approach to regulating hunting of migratory shorebirds can blur its full effects until data are aggregated at a full life-cycle scale (Gallo-Cajiao et al. 2020). Regulatory frameworks for shorebird harvest in particular are variable across the north coast of South America (Watts & Turrin 2016). Shorebirds are totally protected from harvest in Brazil, legally harvested for sport in French Guiana, and entirely unregulated in Guyana, including commercial hunting. This regulatory heterogeneity creates opportunities for laundering, whereby shorebirds illegally harvested in one jurisdiction may be sold in others where regulations are weak or absent. Although international agreements such as the Convention on International Trade in Endangered Species of Wild Fauna and Flora (CITES) and the Convention on Migratory Species (CMS) are relevant, their effectiveness is limited in this context. Suriname, for example, is a party to CITES but not CMS, while migratory shorebird species are listed under CMS but not CITES.

Addressing these challenges will require a coordinated, multi-jurisdictional approach grounded in improved, species-level harvest assessments and population monitoring, allowing harvest levels to be explicitly linked to population trends (Cox et al. 2025). Equally important is a better understanding of the human dimensions of shorebird harvest in the Western Atlantic Flyway, including equity, livelihood dependence, and perceptions of regulations and conservation measures, as policies lacking user acceptance are unlikely to succeed. Both lines of research on harvest effects and human dimensions will need to be further integrated into the policy process balancing conservation and human needs. More specifically, outputs from this research agenda could inform institutional arrangements already in place that have incorporated harvest mitigation as a priority, such as the Atlantic Flyway Shorebird Initiative (AFSI Harvest Working Group 2017, 2020).

## 5. Conclusion

Our study contributes to the understanding of wetland use more generally and to the consumptive use of waterbirds more specifically, both in the tropics as well as in Suriname within the context of central northern South America and the Western Atlantic Flyway. In doing so, we discovered that harvest could be happening at unsustainable levels both for resident and migratory species, supporting global analyses on the importance of overharvesting as a threat to biodiversity (Maxwell et al. 2016). Illegally hunted species are typically those that are of greatest conservation concern. The issue of waterbird harvest, and shorebirds in particular, has been recognized for decades in the central northern coast of South America and the Lesser Antilles (Hutt, 1991). However, structured and systematic assessments have only recently become available, highlighting the challenges involved in addressing this issue. More dedicated resources to monitor the status of populations and the level of harvest will greatly help develop more effective regulatory frameworks for waterbird conservation in the region. However, while enhancing existing regulatory frameworks is important, it is crucial to also prioritize efforts in promoting compliance, engaging with local communities, and developing alternative livelihoods. Given the importance of Suriname for waterbirds in the Americas, developing a cost-effective approach to Suriname’s harvest management of species with international conservation concern, such as the scarlet ibis and migratory shorebirds, should remain a priority for the international community.

## Acknowledgements

We are deeply grateful to all the hunters who participated in this study. We also extend our sincere thanks to Marie Djosetro, Bryan Drakenstein, and Hesdy Esajas for their valuable contributions at the outset of the fieldwork in 2006 and 2016, respectively. Special thanks to Chanderbhan Dwarka, Amir Imraan, Astrid Moor, and Ashok Pherai for their assistance in conducting the surveys, and to Brendan Gray for his support in compiling the hunter database. F. Tremblay, two reviewers, and the editors also provided invaluable comments that greatly helped to improve this manuscript.

## Funding statement

The surveys conducted in 2006 received financial support from NVD Nature Conservation Fund, BirdLife Netherlands. Funding for the survey conducted in 2016 was provided to New Jersey Audubon by US Fish and Wildlife Service, through a grant from National Fish and Wildlife Foundation. The project received additional funding from Prince Bernhard Culture Fund, Dierenrampenfonds Foundation, Van der Hucht De Beukelaar Foundation, and Environment and Climate Change Canada.

## Author contributions

The conception of the project was led by DSM and ALS. The questionnaire was developed by ALS, MJSS, and DSM. Data collection was led by ALS and MJSS. ARC and CR analyzed the data and prepared the tables and figures. All the authors where invovled in the interpretation of the data. The first draft of the paper was written by EGC and ARC. All authors approved the final version of the manuscript.

## Conflict of interest statement

The authors declare no conflicts of interest relevant to this publication.

## Ethic Statement

Informed consent was obtained from all participants before interviews

## Notes

### Competing Interest Statement

The authors have declared no competing interest.

### Summary of Updates

We clarified and strengthened the Results section and streamlined the language in the Discussion. We also identified and corrected a minor data-processing error. Specifically, 72 hunters who had previously been deemed unreliable were inadvertently included in the original analysis due to an omission during data filtering. These individuals have now been excluded, and all analyses have been updated accordingly. Importantly, this correction does not affect the results or the conclusions of the manuscript.

