## Appendix for "A national assessment of waterbird hunting in coastal wetlands of Suriname, South America"

### Appendix 1

**Questionnaire hunting/poaching coastal birds Suriname 2006**

1. In which district are you living?

2. Did you visit the coastal swamps, mangrove forests, or mudflats during the last 12 months? Yes (go to 3)/No (go to 9).

3. If yes, which area did you visit most frequently?

4. What was the aim of your visit to that area? Hunting/Fishing/Recreation/ Others (please, specify).

5. How often did you visit that area during the last 12 months? Once/2-10 times/11-50 times/ more than 50 times.

6. Did you actively hunt in the area during the last 12 months? Yes/No.

7. How many birds of the following species did you shoot in that area during the last 12 months? Indicate 0, 1, some, tens, hundreds, more than one thousand.

Blue-winged Teal, White-cheeked Pintail, Black-bellied Tree-Duck, Muscovy Duck, American Flamingo, Cocoi Heron, Great Egret, various species of small herons, Wood Stork, Jabiru, Scarlet Ibis, Roseate Spoonbill, North-American shorebirds, other species (which ones?)

8 How many birds of the following species are hunted in that area within one year by all hunters in total? Indicate 0, 1, some, tens, hundreds, a few thousand, many thousands.

Blue-winged Teal, White-cheeked Pintail, Black-bellied Tree-Duck, Muscovy Duck, American Flamingo, Cocoi Heron, Great Egret, various species of small herons, Wood Stork, Jabiru, Scarlet Ibis, Roseate Spoonbill, North-American shorebirds, Other species (which ones?)

9. Did you hear, see, or noticed in a different way that during the last 12 months birds of the following species have been shot? Put a cross behind the species name.

American Flamingo, Cocoi Heron, Great Egret, various species of small herons, Wood Stork, Jabiru, Scarlet Ibis, Roseate Spoonbill, North-American shorebirds, Other species (which ones?)

### Appendix 2

**Questionnaire: Hunting coastal birds Suriname 2016**

1. How old are you? ______
   a. Do you have others living in your household? _____. If yes, how many people live in your household?

2. Did you visit the coastal swamps, mangrove forests, or mudflats during the last 12 months? Yes (go to 3)/No (go to 12)

3. Which location did you visit most frequently? ___________________________
   a. What was the purpose of your visit to that location? Hunting/Fishing/ Recreation/Other (please, specify). __________________________________.
   b. How do you access the location (walking, bicycle, motor bike, car, boat)? ____________.

4. How often did you visit that location?
   a. Every day _____. Not every day, but several times per week _____. Once per week _____. 1-3 times per month _____. Not every month, but several times per year _____.

5. Did you hunt in that location during the last 12 months? Yes (go to 6)/No (go to 12). _____.

6. How many birds of the following species did you shoot in that location during the last 12 months? Indicate 0, 1, some tens (10-50) several tens (50-100),some hundreds (100-500), several hundred (500-1000), more than one thousand. Please specify for each.

   Blue-winged Teal, White-cheeked Pintail, Black-bellied Tree-Duck, Muscovy Duck, American Flamingo, Cocoi Heron, Great Egret, various species of small herons, Wood Stork, Jabiru, Scarlet Ibis, Roseate Spoonbill, North-American shorebirds, other species (which ones?)

7. What is your main purpose for hunting? Eat _____. Sell _____. Sport/recreation _____. Other (please specify) ____________________________

8. What is your primary bird hunting method? Shotgun _____. Net _____. Choking wire _____. Other (please specify) ___________________________________.

9. What time of year do you usually hunt? August-November (Long dry season) _____. December-January (Short rainy season) _____. February-March (Short dry season) ______. May-July (Long rainy season) _____.

10. How many birds of the following species do you think are hunted in that location within one year by all hunters in total? Indicate 0, 1, some, tens, hundreds, a few thousand, many thousands. Please specify for each.

    Blue-winged Teal, White-cheeked Pintail, Black-bellied Tree-Duck, Muscovy Duck, American Flamingo, Cocoi Heron, Great Egret, various species of small herons, Wood Stork, Jabiru, Scarlet Ibis, Roseate Spoonbill, North-American shorebirds, Other species (which ones?)

11. Do you hunt more _____, less_____ or the same ____ number of coastal birds as you did 10 years ago? If your answer is "more" or "less," please explain the reasons why.

12. A similar survey to the one you just completed was conducted in 2006. Please indicate if you participated in this survey (Yes/No).

13. Did you hear, see, or noticed in a different way that during the last 12 months birds of the following species have been shot? Put a cross behind the species name.

American Flamingo, Cocoi Heron, Great Egret, various species of small herons, Wood Stork, Jabiru, Scarlet Ibis, Roseate Spoonbill, North-American shorebirds, Other species (which ones?)

END OF QUESTIONAIRRE

### Appendix 3

#### Jags code

```
 model{

#Priors on harvest mean across species and standard deviation for each
for (y in 1:Nyear){
        eta_alpha[y] ~ dnorm(0,pow(5,-2))
       sigma_alpha[y] ~ dt(0,1,4)T(0,)
}

#centered parametrisation trick for random effect
for(j in 1:(Nsp)){
    epsilon_alpha[2,j] ~ dnorm(0,pow(1,-2))
    epsilon_alpha[1,j] ~ dnorm(0,pow(1,-2))
}

#derive mean for each species for fist year
for(j in 1:(Nsp)){
            log(mu_harvest[1,j]) <- eta_alpha[1] + epsilon_alpha[1,j]* sigma_alpha[1]
}

#derive mean for each species for second year
for(j in 1:(Nsp-1)){
            log(mu_harvest[2,j]) <- eta_alpha[2] + epsilon_alpha[2,j]* sigma_alpha[2]
}

#Species 13-14 are large and small shorebirds.
#Their means should sum together to be the mean of the total shorebirds
#(only relevant for 2016)
mu_harvest[2, 15] ~ dsum(mu_harvest[2, 13], mu_harvest[2,14])


#NB2 form (Linden and Mantyniemi 2011)
#size (dispersion) parameter shared across years but unique for each species
for (j in 1:Nsp){
  omega[j] = 1
  theta[j] ~ dgamma(5,0.1)
}

###derive probability and variance from the mean and size parameter
for(y in 1:Nyear){
  for(j in 1:Nsp){
                        sigma_sqr[y,j] <- omega[j]*mu_harvest[y,j] + theta[j] *
pow(mu_harvest[y,j],2)
                        p[y,j] <- mu_harvest[y,j]/sigma_sqr[y,j]
                        r.c[y,j] <-  pow(mu_harvest[y,j],2)/(sigma_sqr[y,j] - mu_harvest[y,j])
}}
```

```
###derive probability for each number of harvest
for(j in 1:Nsp){
  for(y in 1:Nyear){
  cat.prob[y,j,1] <-  dnegbin(0, p[y,j], r.c[y,j])                                        #0
  cat.prob[y,j,2] <-  dnegbin(1, p[y,j], r.c[y,j])                                        #1
  cat.prob[y,j,3] <-  pnegbin(10, p[y,j], r.c[y,j]) - pnegbin(1, p[y,j], r.c[y,j])        #2 to 10
  cat.prob[y,j,4] <-  pnegbin(50, p[y,j], r.c[y,j]) - pnegbin(10, p[y,j], r.c[y,j])       #11 to 50
  cat.prob[y,j,5] <-  pnegbin(100, p[y,j], r.c[y,j]) - pnegbin(50, p[y,j], r.c[y,j])      #51 50 100
  cat.prob[y,j,6] <-  pnegbin(500, p[y,j], r.c[y,j]) - pnegbin(100, p[y,j], r.c[y,j])     #101 to 500
  cat.prob[y,j,7] <-  pnegbin(1000, p[y,j], r.c[y,j]) - pnegbin(500, p[y,j], r.c[y,j])    #500 to 1000
  cat.prob[y,j,8] <-  1 - pnegbin(1000, p[y,j], r.c[y,j])                                 # > 1000
}}

#Harvest likelihood-- this is only for active harvesters.
for (i in 1:Nresp){
  for (j in 1:Nsp){
    Harvest[i, j] ~ dcat(cat.prob[YearIndex[i],j, 1:Nbins])
}}


#Fit statistics
for ( i in 1:Nresp){
  for (j in 1:Nsp){
    Harvest_new[i, j] ~ dcat(cat.prob[YearIndex[i],j, 1:Nbins])
    ld_harv_new[i,j] <- logdensity.cat(Harvest_new[i, j], cat.prob[YearIndex[i],j, 1:Nbins]) *
fitmask[i,j]
    ld_harv[i,j] <-  logdensity.cat(Harvest[i, j], cat.prob[YearIndex[i],j, 1:Nbins]) * fitmask[i,j]
}}

fit_harv <- sum(-2*ld_harv)
fit_harv_new <- sum(-2*ld_harv_new)

#end of model
}
```
